# A minimum effective dose for (transcranial) alternating current stimulation

**DOI:** 10.1101/2022.07.06.499054

**Authors:** Ivan Alekseichuk, Miles Wischnewski, Alexander Opitz

**Author notes:** Corresponding author. Dept of Biomedical Engineering, University of Minnesota, 312 Church St. SE, 7-105 Nils Hasselmo Hall, Minneapolis, MN 55455, USA. E-mail address (I. Alekseichuk).

## Abstract

Transcranial alternating current stimulation (tACS) is a popular method for non-invasive neuromodulation in human research and therapy. However, what constitutes an effective dose in tACS applications is still a matter of investigation and debate. Here, we examined available literature data regarding the effects of alternating current (AC)-induced electric fields on cellular-level neural activity. The literature search identified 16 relevant experimental reports that utilized brain slices, anesthetized rodents, and awake/behaving animal models. We implemented a probabilistic meta-analysis to estimate the minimum effective dose (MED) required for inducing detectable significant neural changes with AC stimulation. The results showed that AC stimulation at 0.3 mV/mm in awake/behaving mammals leads to an 80% probability of inducing minimum neural effects. A similar level of effectiveness in brain slices and anesthetized mammals required 0.7 mV/mm. In conclusion, (transcranial) alternating current stimulation is significantly more effective in awake than in anesthetized brains. The proposed dose targets can serve as a practical guideline for tACS in humans.

## Introduction

The question of adequate dosing is fundamental to non-invasive brain stimulation. Selecting stimulation intensity that results in an electric field leading to a desired physiological response is crucial. For transcranial alternating current stimulation (tACS), the concept of dose refers to the induced electric fields in the brain [1–3]. Thus, we can ask: What minimum electric field strength causes detectable changes in neural activity? This question is critical for understanding tACS dosing in humans necessary to modulate mental states and behavior. Many in-vitro and animal studies have investigated how intrinsic and induced oscillating electric fields modulate neural activity. However, a clear consensus on an adequate electric field strength for physiological effects is still lacking. Here, we use a meta-analytic probabilistic approach to determine the minimum effective dose (MED) at which alternating current (AC) modulates neural spiking.

## Methods

To identify effective stimulation parameters, we manually collected data from 16 papers identified in a PubMed® search (see Supplementary Table S1). We included experimental mechanistic studies in brain slices and animals that applied AC stimulation at 0.1-100 Hz with simultaneous direct neural recordings. In addition, we only considered papers that measured the AC electric field. Our analysis summarized 11 experiments in rodent brain slices and deeply anesthetized rodents (study group I) and 5 experiments in awake or behaving nonhuman primates and rodents (group II). From them, we extracted the lowest electric field strength values at which neural changes were detected, which constitutes the upper MED boundary (a certainly effective dose per experiment). The highest electric field values at which no changes occurred defined the lower MED boundary (a certainly ineffective dose per experiment). Here we define the “minimum effect” as significant changes in at least one observed unit in the sample. See Supplementary Methods for more details. We assumed that the MED lies between these boundaries and follows a Beta distribution, commonly used in uncertainty analyses. The meta-analytical distribution is a Gaussian function, satisfying the Central Limit Theorem, truncated to avoid negative values. Moreover, we estimated 99% confidence intervals (CI_99_) of the meta-analytical fit, assuming that each underlying study has a statistical power of 80%. Finally, study groups were statistically compared using the nonparametric two-sample Kolmogorov-Smirnov (KS) test.

**Fig. 1.**
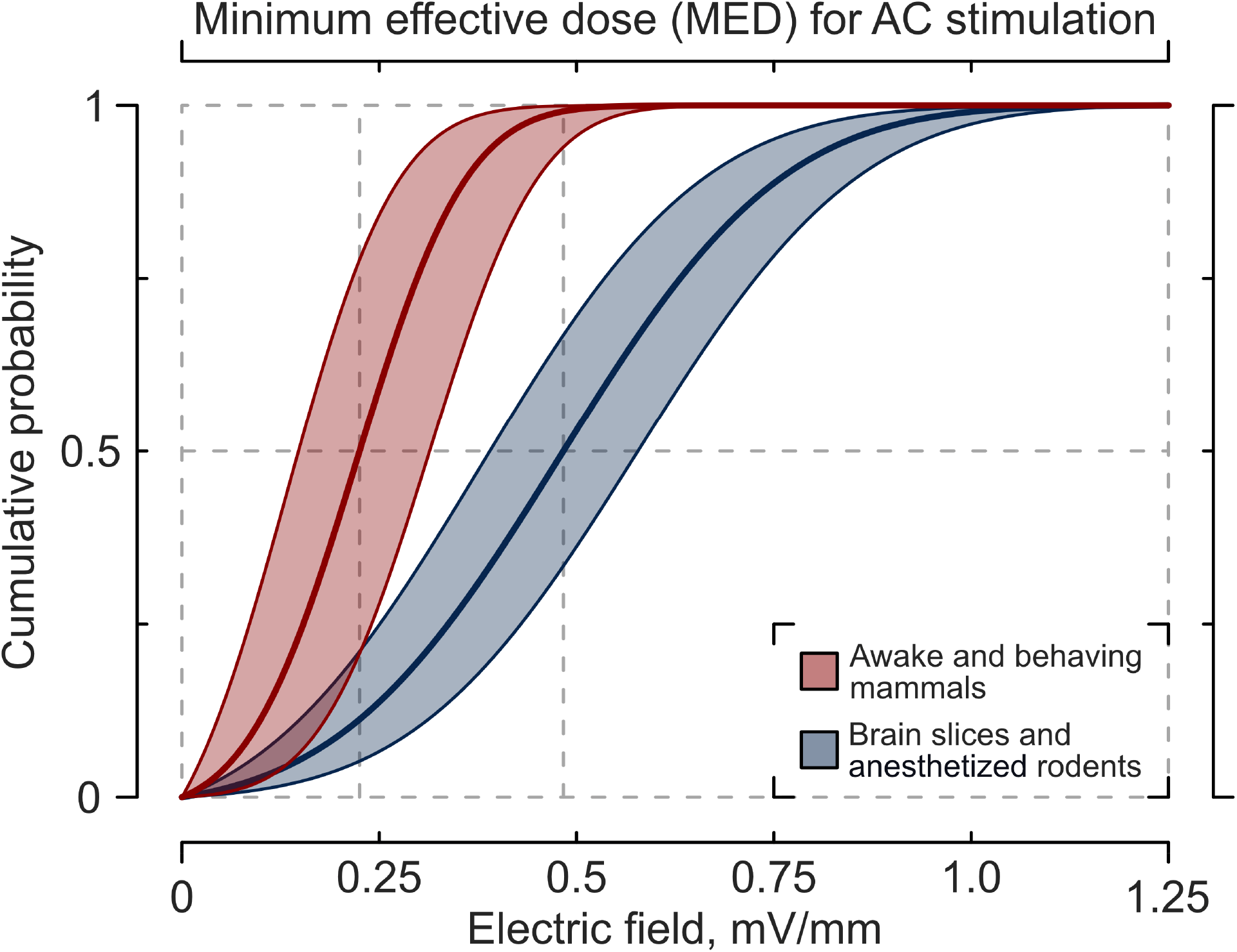
Meta-analysis of alternating current (AC)-induced electric field magnitudes that led to minimum significant neural changes. The probability functions were fitted into the literature data from studies in brain slices and anesthetized rodents (blue) and awake/behaving rodents and nonhuman primates (red). The shaded areas correspond to CI_99_.

## Results and Discussion

Meta-analytic results show the mean probabilistic MED of 0.49 mV/mm (CI_99_: 0.39-0.58 mV/mm) and standard deviation of 0.21 mV/mm for AC stimulation in brain slices and anesthetized rodents. This mean dose corresponds to the cumulative probability of 50% to elicit minimum significant neural changes. Results in brain slices and anesthetized animals separately are given in Supplementary Materials (Fig. S1). For awake/behaving mammals, the MED’s mean = 0.23 mV/mm (CI_99_: 0.15-0.31 mV/mm) and SD = 0.10 mV/mm. By extension, the 80% probability of eliciting a significant neural response corresponds to 0.67 mV/mm (CI_99_: 0.57-0.76 mV/mm) in the slices and anesthetized rodents and 0.31 mV/mm (CI_99_: 0.23-0.40 mV/mm) in awake/behaving mammals. The required dose for the latter is 2.16 times lower than for the former, which is significant according to the KS test (p-value < 0.001). We should notice that the awake brain activity can spontaneously fluctuate, enabling accidental alignment to tACS over time. Thus, the comparison of “awake” to “fixed” brains should be taken with some caution.

Our analysis indicates that sub-millivolt per millimeter oscillating electric fields can elicit measurable neural modulation. Such electric field strengths can be achieved in humans using tACS at 2-4 mA [2,3] while mitigating tolerability issues with distributed multi-electrode stimulation montages. Crucially, the effective dose threshold here is twice lower for an active brain than for a “fixed” brain, highlighting a major role of intrinsic neural states. This conclusion agrees with individual studies. Ozen et al. demonstrated that AC stimulation at ~0.67-1 mV/mm (recalculated from the reported 0.8-1.2 V) induced a response in rats under anesthesia, while ~0.33 mV/mm was already sufficient to modulate their brain during normal sleep [4]. Conversely, even ~1.67 mV/mm had no effects in rats during exploratory behavior. Furthermore, Fröhlich and McCormick showed that stimulating brain slices with their “natural” AC waveform (recorded and played back) makes the neurons twice more susceptible to entrainment in comparison to regular AC stimulation [5]. Huang et al. stimulated posterior alpha oscillations in ferrets and showed better neural susceptibility to tACS closer to intrinsic frequencies [6]. On the other hand, Johnson and Alekseichuk et al. found no straightforward relationships between the spiking rates of individual neurons and their response to a given stimulation frequency [7]. One important factor is that all original studies here investigated the immediate neural response due to short stimulation, leaving the question of long-lasting changes outside the scope. Noteworthy, the stimulation frequency itself can play a particular role. Several studies reported that stimulation at lower frequencies (<20 Hz) is more effective than at higher frequencies (50-100 Hz) [1,8,9]. Finally, different brain regions can have different innate thresholds to AC stimulation due to their cytoarchitecture. Francis et al. found a twice lower MED in CA1 vs. CA3 rat slices, all other factors equal [10].

Our analysis addresses the minimum effective dose, but physiological responses at above threshold intensities are still under active investigation. Recently, Krause et al. reported that neurons intrinsically entrained to a particular oscillation can be desynchronized by tACS at the same frequency for low intensities before becoming re-entrained at higher intensities [11]. Here, we analyzed the studies exploring the tACS effects on neural activity to find the minimum effective dose. Nevertheless, the exact way to quantify neural activity varies between studies, adding variability to the analysis. Crucially, “how much?” (dose) is not the only stimulation factor that defines the response; the others are “where?” (location) and “what?” to stimulate (frequency). A full dose-response curve and its dependency on every factor are beyond the scope of the present analysis, as the existing studies explored different parameters while extensive parametric experimental studies are yet to be performed. A given experimental study could consider reaching for an above-minimum target dose when stronger effects are required.

In summary, we show that AC electric fields as low as 0.3 mV/mm can be sufficient to modulate neurons in awake and behaving mammals and can thus inform one of the minimum requirements for tACS dosing. Notably, these magnitudes are twice lower than required for anesthetized rodents and brain slices, and future research is needed to determine if task engagement in humans further increases the brain’s susceptibility to tACS.

## Acknowledgments

This work was supported by the Brain & Behavior Research Foundation (Young Investigator Grant to I.A.), the University of Minnesota’s MnDrive Initiative (M.W. and A.O.), and the National Institutes of Health (grant RF1MH124909 to A.O.).

### Appendix A. Supplementary data

Supplementary methods and data to this article can be found online.

