## Supplementary Materials for "A minimum effective dose for (transcranial) alternating current stimulation"

### Supplementary Methods

In the present analysis, we define the "dose" as a quantity of stimulation ("how much?") in terms of the magnitude of the induced electric field (mV/mm). Other crucial factors of alternating current (AC) stimulation, such as electrode montage ("where" to stimulate) and frequency ("how" to stimulate), remain outside the scope of the present work. We define "minimum effect" as detectable, significant changes in at least one unit of observation (e.g., neuron) per sample (e.g., brain). Observable changes in neural activity could have occurred as significant differences in the number of neural spikes per time (firing rate) and/or spike timing (phase entrainment). Because each study observed a highly limited number of units (e.g., neurons) relative to the whole sample (e.g., brain), significant changes in a few units are sufficient to presume potential impact on the whole sample. Our confidence intervals estimation of the meta-analysis below further mitigates the putative impact of outliers here.

To extract the data for meta-analysis, we first identified principal experiments per each original paper. Those experimental conditions were performed using different cell tissue types, stimulation waveforms, and behavioral conditions (if available). Alternating current frequency was not considered an independent factor and aggregated in the meta-analysis. This is because there is little agreement between the studies on exact applied frequencies and the moderate number of studies. Thus, it is not feasible to estimate the minimum effective dose per frequency or confirm the existence of such frequency-dose relationships. At the same time, all utilized stimulation frequencies are in the same range, typical for tACS in humans (<100 Hz), allowing their aggregation. From each principal experiment, we extracted up to two data points. First is the lower boundary – the lowest electric field magnitude at which the effects were observed. Second is the upper boundary (if available) – the highest ineffective electric field magnitude given for the same stimulation frequency and other parameters as for the lower boundary. Whenever multiple study samples (slices, animals) were examined per principal experiment or the same sample was measured in different locations simultaneously, we aggregated these data.

The electric field magnitudes are taken into the meta-analysis according to the measurements stated in the original papers. It is plausible that in some studies, the recording electrodes were not perfectly oriented to capture these values; however, we assumed a random error in electric field measurements between the studies, which will be partially averaged out during the meta-analysis.

We assumed that the MED lies between the lower and upper boundaries and follows a Beta distribution (see Fig. S2), commonly used in uncertainty analyses. We implemented a symmetric bell-shaped $Beta_{(3,3)}$ function for studies with known boundaries. The lower boundary became zero in experiments with no measurements at a lower-than-effective dose. There, we used a more conservative, right-leaning $Beta_{(4,2)}$ function to deemphasize close-to-zero values. The functions were fitted by minimizing their negative loglikelihood (see Fig. S3). Moreover, we estimated 99% confidence intervals ($CI_{99}$) of the meta-analytical fit using a bootstrapping method (10,000 iterations), assuming that each underlying study has a statistical power of 80% (see Fig. S4).

### Supplementary Tables

**Table S1.** Summary of identified studies exploring the direct neural effects of alternating currents (AC) and AC-induced electric fields. Experiments using different cell tissue types, stimulation waveforms, and behavioral conditions are separate entrances. The lowest electric field magnitude (in mV/mm throughout the manuscript) induced minimum significant neural changes is taken as the upper boundary of the minimum effective dose (MED). The highest electric field magnitude that was ineffective is a lower boundary of the MED (if known, otherwise presumed to be zero). Note that a parallel-to-slice electrode montage for slice studies means that the electric current and electric field were perpendicular to the somato-dendritic axis of pyramidal neurons, which is the most optimal direction.

| Paper | PMID | Object | Electrode montage | Stim. duration | AC waveform | Lower MED boundary | Upper MED boundary |
|---|---|---|---|---|---|---|---|
| Study group I. Brain slices and deeply anesthetized rodents | | | | | | | |
| Bawin et al., 1984; Bawin et al., 1986 | 6098340; 3942883 | CA1 slices (rat) | Parallel to slice | 5-30 s | 5 and 60 Hz | - | **0.7** |

| | | | | | | | |
|---|---|---|---|---|---|---|---|
| Francis et al., 2003 | 12917358 | CA3 slices (rat) | Parallel to slice | ≈ 0.1 s | Gauss pulse 0.5-1 Hz | **-** | **0.787** |
| | | CA1 slices (rat) | Parallel to slice | ≈ 0.1 s | Gauss pulse 1-2 Hz | **-** | **0.298** |
| Deans et al., 2007 | 17599962 | CA3 slices (rat) | Parallel to slice | 1 s | 50 Hz | **-** | **0.25** |
| Radman et al., 2007 | 17360926 | CA1 slices (rat) | Parallel to slice | n/a | 30 Hz | **0.5** | **1** |
| Frohlich & McCormick, 2010 | 20624597 | Visual cortex slices (ferret) | Parallel to slice | 60 s | 0.075–0.375 Hz | **0.5** | **1** |
| | | | | | Intrinsic LFP replayed | **0.25** | **0.5** |
| Reato et al., 2010 | 21068312 | CA3 slices (rat) | Parallel to slice | 2 s | 7 Hz | **-** | **0.5** |
| Ozen et al., 2010 | 20739569 | Rats under anesthesia | A pair on temporal bones – calvarium midline | 15-60 cycles | 0.8-1.7 Hz | **0.333** | **0.667** |
| Anastassiou et al., 2011 | 21240273 | S1 slices (rat) | Parallel to slice | 5 s | 1 Hz | **-** | **0.74** |
| Asamoah et al., 2019 | 30655523 | Rats under anesthesia | Anterior – posterior to the motor cortex | 60 s | 1-2.5 Hz | **0.5** | **1** |
| Study group II. Awake/behaving rodents and nonhuman primates (NHP) | | | | | | | |
| Ozen et al., 2010 | 20739569 | Behaving rats | A pair on temporal bones – calvarium midline | 15-60 cycles | 1.25 Hz | **-** | **0.333** |
| Kar et al., 2017 | 28137971 | Behaving NHP | Anterior central – behind the area MT | 3 s per task trial | 10 Hz | **-** | **0.12** |
| Krause et al., 2019; Krause et al., 2022 | 30833389; 35613140 | Awake NHP | Frontal – occipital electrodes | 300 s | 5-20 Hz | **-** | **0.28** |
| Johnson & Alekseichuk et al., 2020 | 32917605 | Awake NHP | Bilateral temporal electrodes | 180 s | 10 Hz | **-** | **0.38** |
| Huang et al., 2021 | 34035240 | Awake ferrets | Left frontal – neck electrodes | 90 s | Individual alpha rate | **0.25** | **0.5** |

**Supplementary Figures**

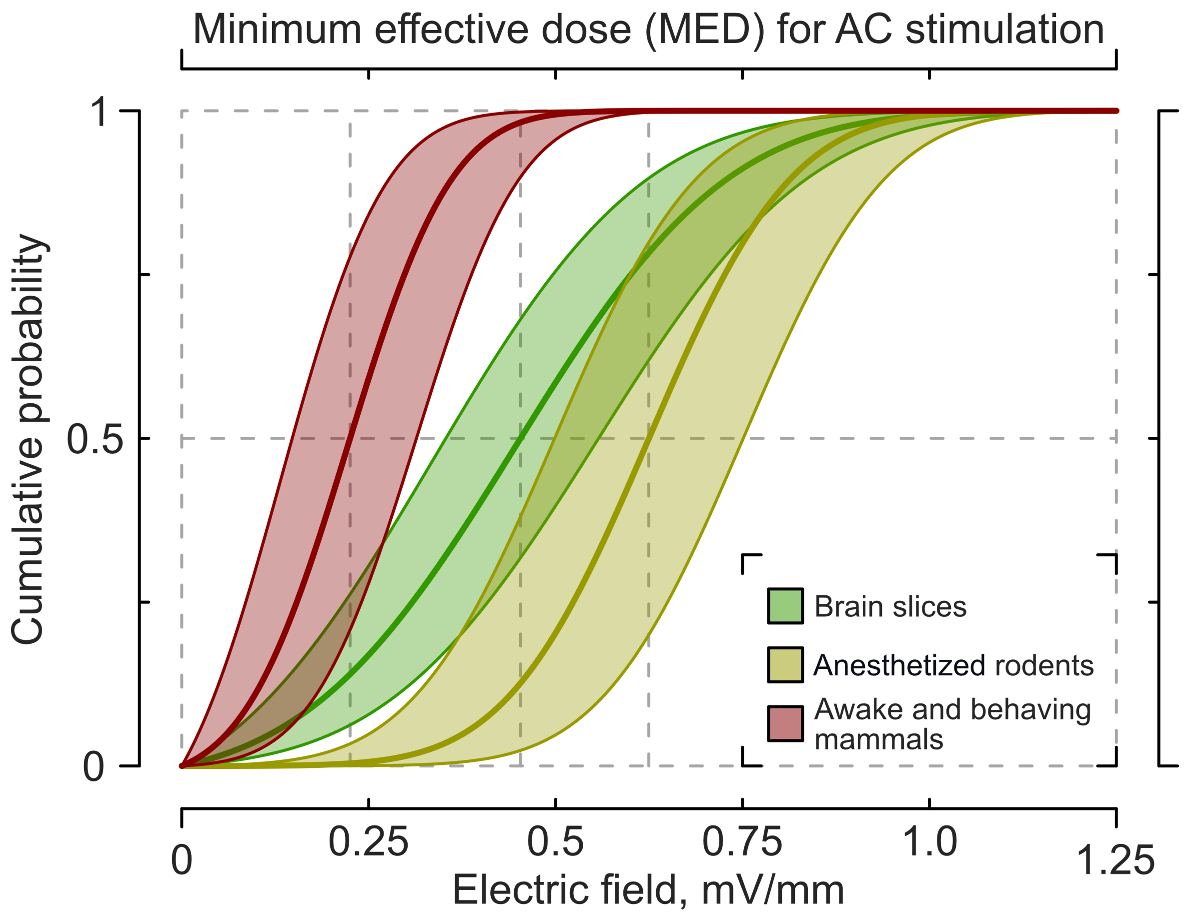


**Fig. S1.** Supplementary meta-analysis related to Fig. 1 in the main text. The experimental studies are regrouped into three categories: brain slices only (n = 9, green), anesthetized rodents only (n = 2, yellow), and awake/behaving rodents and nonhuman primates (n = 5, red). The shaded areas correspond to 99% confidence intervals of the meta-analytical fit. The doses that have an 80% probability to elicit a minimum effect are 0.64 [0.53-0.74] V/mm for the brain slices, 0.75 [0.63-0.88] mV/mm for the anesthetized rodents, and 0.31 [0.23-0.40] mV/mm for the awake/behaving mammals.

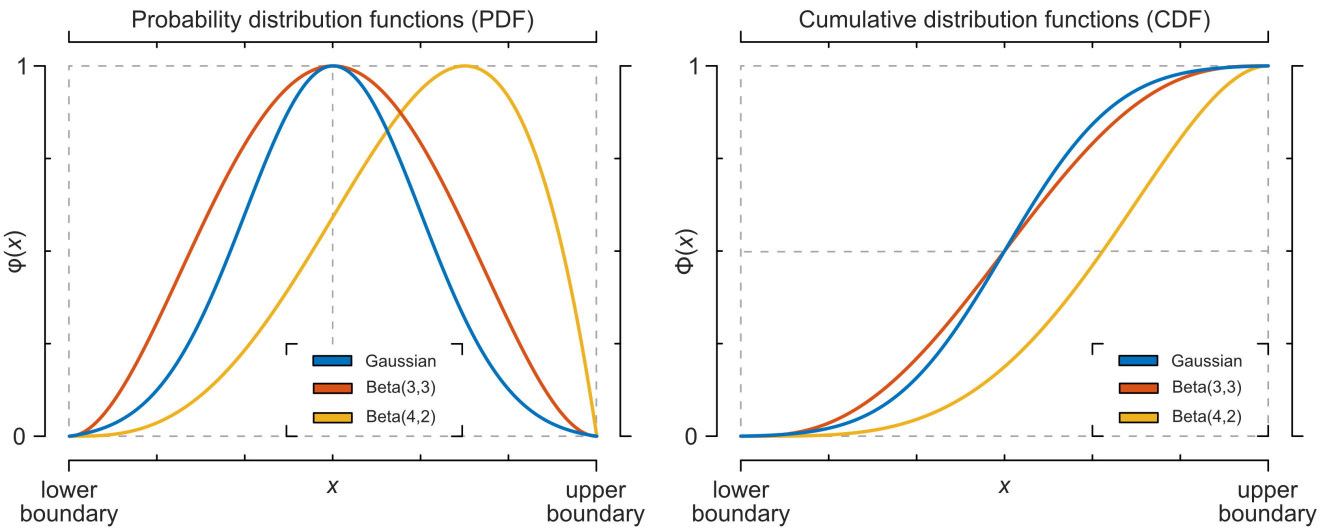


**Fig. S2.** Probability distribution functions (PDF) and cumulative distribution functions (CDF) of truncated Gaussian or Normal distribution (blue), symmetric bell-shaped $Beta_{(3,3)}$ distribution (orange), and right-leaning $Beta_{(4,2)}$ distribution (yellow). The functions are rescaled to their lower and upper boundaries (for Beta from 0 to 1; for Gaussian from $-3\sigma^2$ to $+3\sigma^2$) for illustrative purposes.

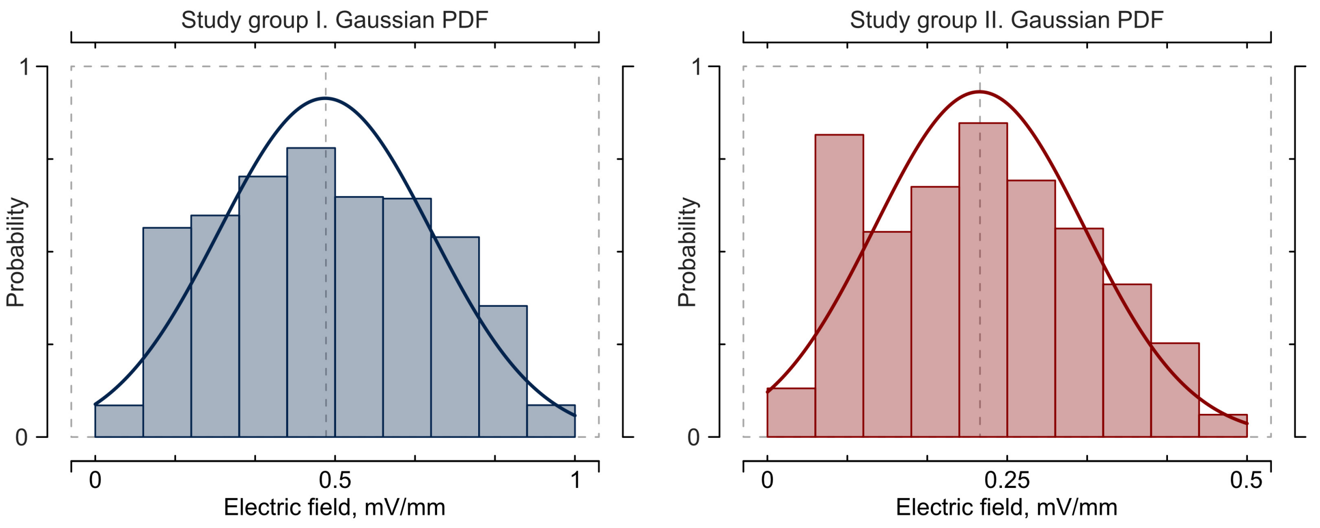


**Fig. S3.** Summary of the minimum effective dose (MED) estimated from the studies in Table S1 and their Gaussian probability density functions (PDFs). Data from studies in brain slices and anesthetized rodents (Group I) are depicted in blue and from awake/behaving rodents and nonhuman primates (Group II) – in red.

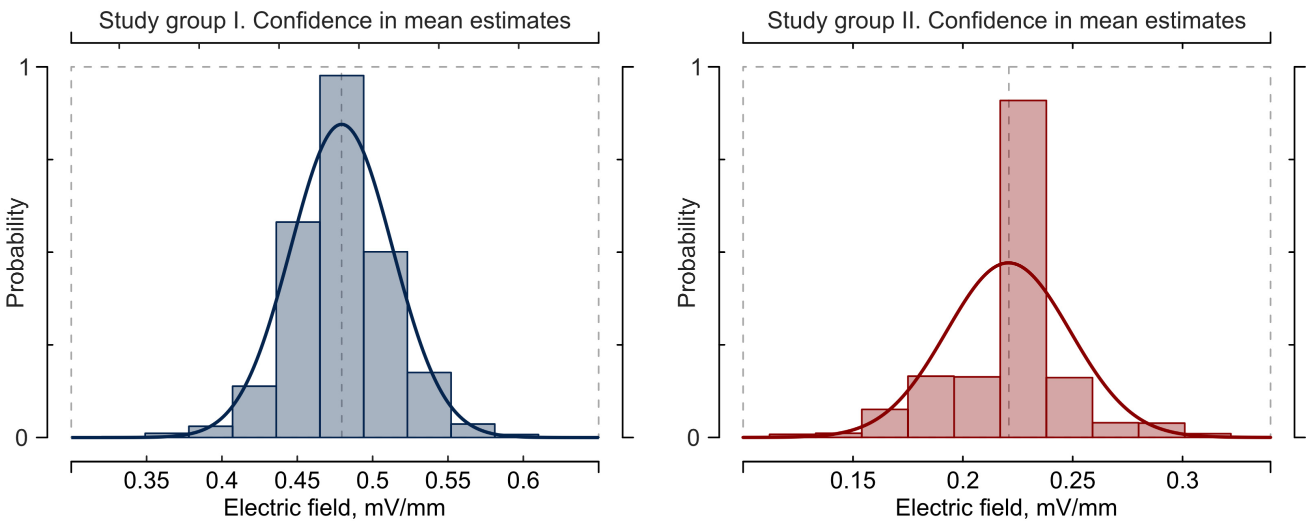


**Fig. S4.** The mean value estimates during the meta-analytical Gaussian fit. The distributions are computed using a bootstrapping method (10,000 iterations), assuming that each underlying study has a statistical power of 80%. Data from studies in brain slices and anesthetized rodents depicted in blue and awake/behaving rodents and nonhuman primates – in red.
